# Amoxicillin inactivation by thiol-catalyzed cyclization reduces protein haptenation and antibacterial potency

**DOI:** 10.1101/647966

**Authors:** María A. Pajares, Tahl Zimmerman, Francisco J. Sánchez-Gómez, Adriana Ariza, María J. Torres, Miguel Blanca, F. Javier Cañada, María I. Montañez, Dolores Pérez-Sala

## Abstract

Serum and cellular proteins are targets for the formation of adducts with the β-lactam antibiotic amoxicillin. This process could be important for the development of adverse, and in particular, allergic reactions to this antibiotic. In studies exploring protein haptenation by amoxicillin, we observed that reducing agents influenced the extent of amoxicillin-protein adducts formation. Consequently, we show that thiol-containing compounds, including dithiothreitol, N-acetyl-L-cysteine and glutathione, perform a nucleophilic attack on the amoxicillin molecule that is followed by an internal rearrangement leading to amoxicillin diketopiperazine, a known amoxicillin metabolite with residual activity. The effect of thiols is catalytic and can render complete amoxicillin conversion. Interestingly, this process is dependent on the presence of an amino group in the antibiotic lateral chain, as in amoxicillin and ampicillin. Furthermore, it does not occur for other β-lactam antibiotics, including cefaclor or benzylpenicillin. Biological consequences of thiol-mediated amoxicillin transformation are exemplified by a reduced bacteriostatic action and a lower capacity of thiol-treated amoxicillin to form protein adducts. Finally, modulation of the intracellular redox status through inhibition of glutathione synthesis influenced the extent of amoxicillin adduct formation with cellular proteins. These results open novel perspectives for the understanding of amoxicillin metabolism and actions, including the formation of adducts involved in allergic reactions.

## Introduction

Allergic reactions to drugs and to antibiotics in particular, constitute important medical problems [1, 2]. These reactions can be life-threatening and limit the therapeutic tools against infections. Binding of β-lactam antibiotics to endogenous protein carriers is considered an important step in the development of allergic reactions [3]. Nevertheless, our knowledge of this process is still limited. It is generally accepted that the low molecular weight of antibiotics and other drugs prevents them from being recognized by the immune system on their own, for which they require association with larger structures [4]. Importantly, numerous drugs or their metabolites are able to bind covalently to proteins, a process known as haptenation [5, 6], which renders structures with the potential of being recognized by the immune system and induce an allergic response [7] (Fig. 1).

**Fig. 1.**
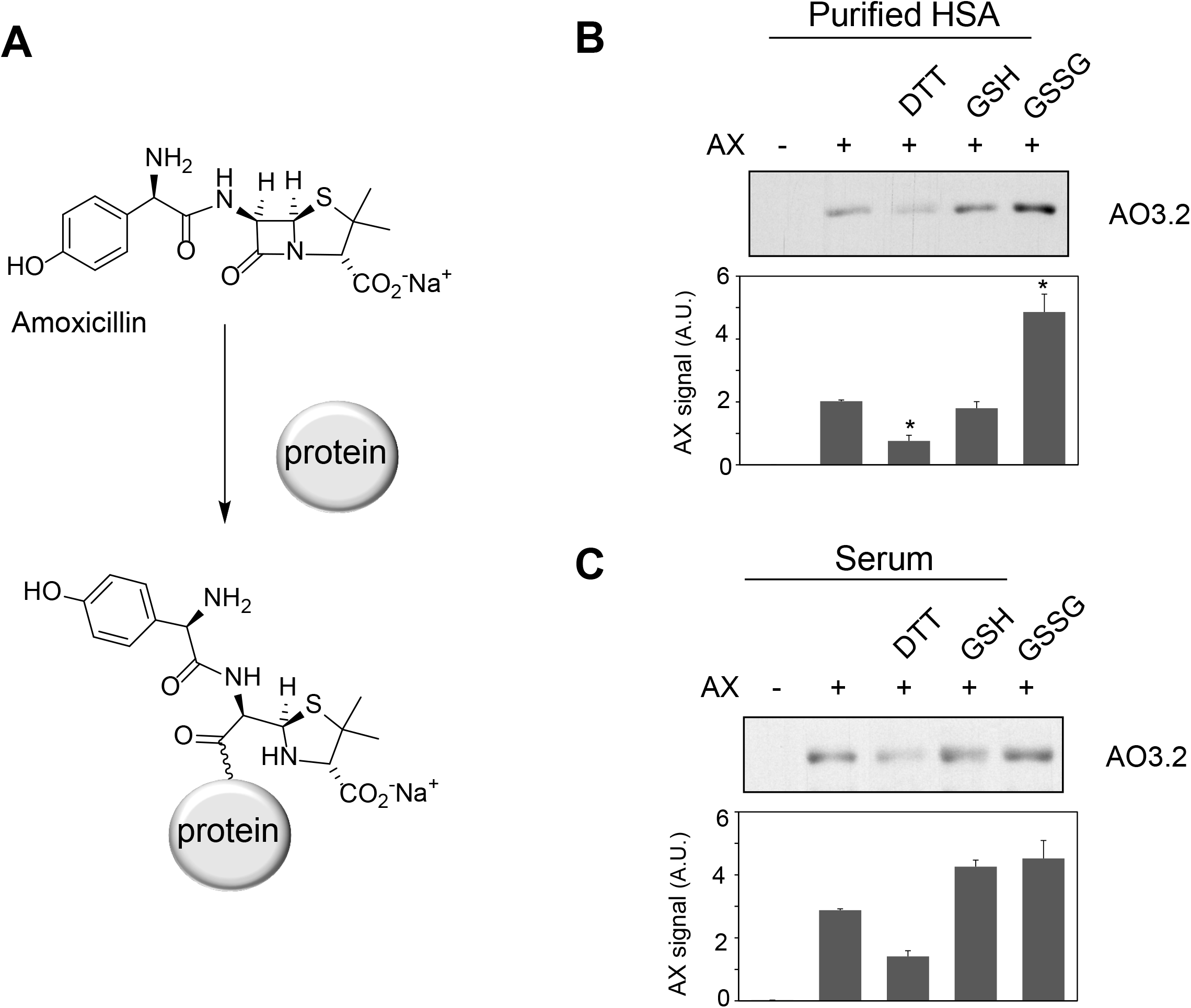
Effect of various compounds on amoxicillin-protein adduct formation. **(A)** Scheme of the formation of amoxicillin-protein adducts. Purified HSA **(B)** at 10 mg/ml in PBS or total human serum **(C)** was preincubated with vehicle, 1 mM DTT, 2.5 mM GSH or GSSG for 6 h at 37°C, after which, amoxicillin (AX) from a stock solution in bicarbonate buffer was added at a final concentration of 0.5 mg/ml, as indicated, and incubation was continued overnight at 37°C. Aliquots from the incubation mixtures containing 2 μg of protein were separated by SDS-PAGE. The formation of AX-HSA adducts was monitored by immunoblot with monoclonal anti-amoxicillin AO3.2 antibody. Band intensity was estimated by image scanning and average results ± SEM from three independent assays are shown in the corresponding graphs (*p≤0.05).

Recent work of our group identified several serum proteins as potential targets for haptenation by amoxicillin (AX) [8]. In addition, intracellular proteins are also susceptible of covalent modification by this antibiotic [9, 10]. Nucleophilicity and a favorable environment appear to be important for the selective modification of proteins on certain residues. In vitro, incubation of human serum albumin (HSA) with AX, results in the modification of several lysine residues of which, Lys190 and Lys195 appear to be the most reactive [8]. Interestingly, after oral administration of AX, only modified Lys190 has been found [11, 12]. Apart from the chemical reactivity of the antibiotic and the protein residues, little is known about other factors influencing protein haptenation by AX in biological settings.

Previous studies with the antibiotic sulfamethoxazole have shown that oxidative stress can potentiate the formation of protein adducts in cells [13]. This alteration of the redox status concurs with many diseases and is involved in their pathogenesis or perpetuation. Therefore, oxidative stress is a concomitant factor in numerous pathological situations requiring antibiotic administration. Oxidative stress can influence drug protein-adduct formation by multiple mechanisms, including induction of conformational alterations or oxidative modifications that can either increase or blunt the reactivity of the modified residues towards the drug, as we have recently reviewed [14]. While exploring the impact of oxidative or reducing agents on protein modification by AX, we observed that the presence of exogenous thiol-containing molecules influenced the extent of AX-protein adduct formation. Investigation into the basis of this effect has unveiled the capability of thiol-containing compounds to catalytically transform AX. These results shed new light into the factors determining AX bioavailability and, therefore, its antimicrobial potency and adduct formation capacity, with potential implications for the understanding of drug allergy.

## Materials and Methods

### Reagents

Dithiothreitol (DTT), glutathione (GSH), 2-mercaptoethanol, β-NADPH, 5,5-dithio-bis-(2-nitrobenzoic acid), ethanothiol, triethanolamine, 2-vinylpyridine, sulfosalicylic acid, human serum albumin (HSA), bovine serum albumin (BSA), buthionine sulfoximine (BSO), tris(2-carboxyethyl)phosphine (TCEP), cefaclor and cephadroxil were products of Sigma. Glutathione reductase (GSR) and Triton X-100 were purchased from Roche and Calbiochem, respectively. Microcon cells and Immobilon-P Transfer Membrane were obtained from Amicon and Millipore, respectively. The bicinchoninic acid (BCA) kit for protein determination was from Pierce. Amoxicillin (Clamoxil) was from Beecham. Ampicillin (AMP), benzylpenicillin (BP) and cefuroxime were from Laboratorios Normon (Madrid, Spain). Deuterium oxide (D_2_O) for proton-nuclear magnetic resonance (^1^H-NMR) studies was from VWR. ECL was purchased from GE Healthcare.

### Synthesis and characterization of diketopiperazine derived from AX

Sodium AX (19 mg, 0.05 mmol) and DTT (15 mg, 0.1 mmol) were dissolved in 1.5 mL of degassed H_2_O under N_2_ atmosphere and stirred overnight at room temperature. The mixture was lyophilized to yield a white solid containing diketopiperazine (DKP) as well as oxidized and reduced forms of DTT. The resulting crude was analyzed by ^1^H-NMR, attenuated total reflection-Fourier transform infrared (ATR-FTIR) spectroscopy and electrospray ionization-mass spectrometry (ESI-MS). The conversion of AX to DKP was quantitative.

^1^H-NMR spectra were recorded at 400 MHz using a NMR Bruker Ascend™ 400 MHz for ^1^H, and D_2_O or deuterated-phosphate buffer saline (PBS-d) as solvents. ^1^H-NMR signals corresponding to DKP were the following: ^1^H-NMR (400 MHz, D_2_O) δ ppm: 7.18 (d, 2H, *J*=8.1Hz), 6.83 (d, 2H, *J*=8.1Hz), 5.19 (s, 1H, CHPh), 5.16 (d, 1H, *J*=2.6Hz, H-6), 4.02 (d, 1H, *J*=2.6 Hz, H-5), 3.37 (s, 1H, H-3), 1.52 (s, 3H, CH_3_), 1.15(s, 3H, CH_3_).

ATR-FTIR spectra were recorded with a FTIR spectrometer (VERTEX 70) equipped with a Golden Gate Single Reflection Diamond ATR System from Specac.

Mass spectra were acquired on a High Resolution Mass Spectrometer Orbitrap, Q-Exactive (Thermo Fisher Scientific) equipped with electrospray ionization (H-ESI-II), coupled to a Liquid Chromatographer Dionex Ultimate 3000 HPLC system (Thermo Fisher Scientific). Solid samples were dissolved in MilliQ water for the analysis of AX and in a MilliQ water-methanol mixture for the analysis of the resulting product (DKP) [15]. Detection was carried out with positive polarity, in selected ion monitoring (SIM mode), selecting 366 ions and MS/MS mode for confirmation of analytes. ESI-MS *m/z* (%) were as follows: 366 (M^+^, 20), 207 (16), 160 (100).

### *In vitro* incubations of β-lactam antibiotics for ^1^H-NMR studies

Incubations of AX with DTT, other thiol-containing compounds or antioxidants were carried out at 37°C and physiological pH in PBS using different ratios of reactants and times. The resulting species were compared with a reference control solution containing only AX kept under the same conditions, following two methodologies: (i) after incubation in PBS in inert atmosphere, mixtures were lyophilized and the resulting solids were dissolved in D_2_O and analyzed by ^1^H-NMR; and (ii) incubations were monitored directly in deuterated-PBS by ^1^H-NMR at different times. DKP NMR signals in PBS (400 MHz, PBS-d) δ ppm: 7.27 (d, 2H, *J*=8.6Hz), 6.93 (d, 2H, *J*=8.6Hz), 5.28 (s, 1H, CHPh), 5.26 (d, 1H, *J*=2.6Hz, H-6), 4.12 (d, 1H, *J*=2.6 Hz, H-5), 3.47 (s, 1H, H-3), 1.62 (s, 3H, CH_3_), 1.26(s, 3H, CH_3_).

Incubations of other β-lactams (BP, AMP, cefaclor, cephadroxil or cefuroxime; 0.05 mmol) with DTT (0.1 mmol) at 37°C were carried out either at physiological (PBS-d, 0.6 mL) or at basic pH (Na_2_CO_3_ in D_2_O, pD=11, 0.6 mL), and compared with a reference control solution containing the β-lactam under the same conditions of pH and temperature. Solutions were monitored directly by ^1^H-NMR at different times.

### Formation of AX-HSA adducts

Mixtures of AX (5 mg/ml) with vehicle or antioxidants (1 mM TCEP, 1 mM Trolox or 100 mM DTT in PBS, pH 7.4) were preincubated for 24 h at 37°C or prepared fresh. Both types of mixtures were diluted 1:1 (v/v) with HSA at 0.6 mg/ml in PBS and further incubated for 24 h more at 37°C, after which, aliquots containing 4 μg of protein were loaded on 12% SDS-PAGE gels and analyzed by western blot.

### Colony formation

Mixtures of AX preincubated with vehicle or the reducing agents as above, were diluted with LB agar to prepare plates with a theoretical antibiotic concentration of 100 μg/ml. Equivalent volumes of PBS were added to control plates. *E. coli* DH5α were seeded on the plates with the resulting mixtures and colony formation was assessed by visual inspection and counting after overnight incubation at 37°C.

### Bacterial growth curves

The antibiotic (137 mM) was incubated with the selected thiol reagents (DTT, GSH, 2-mercaptoethanol) at several molar ratios (1:0 up to 1:20) for 24 hours at 37°C in a final volume of 43 μl in PBS. In parallel, *E. coli* DH5α were grown on LB medium (10 mL) overnight at 37°C and the A_620_ of the culture measured before the experiment. Putative AX inactivation was then evaluated following *E. coli* growth in the presence of the treated antibiotic using multi-well plates. For this purpose, appropriate volumes of the AX inactivation or control reactions were added to each well to obtain a theoretical antibiotic concentration of 75 μg/ml in a final volume of 50 μl. The growth curve (150 μl/well) started by addition of 100 μl of the diluted bacterial culture (A_620_ ~0.1) and was followed by A_620_ at 20 min intervals for 350 min. Measurements were carried out in triplicate for each condition and bacterial growth analyzed using GraphPad Prism software.

### Cell culture

Human lymphocyte RPMI 8866 cells were grown in DMEM supplemented with 10% (v/v) fetal bovine serum (FBS, PAA Laboratories). Cells (10^6^) were treated with vehicle or BSO (50 or 100 μM) for 24 h in complete medium without antibiotics, after which, cells were washed with serum free medium and incubated in this medium in the absence or presence of BSO or AX (2.5 mg/ml) for 16 h. Cells were harvested by centrifugation and washed with PBS before lysis using 10 mM Tris/HCl pH 7.5 containing 0.1 mM EDTA, 0.1 mM EGTA, 0.1 mM 2-mercaptoethanol, 320 μg/ml Pefablock, 2 μg/ml trypsin inhibitor, 2 μg/ml aprotinin, 2 μg/ml leupeptin, 0.1 mM sodium orthovanadate, 50 mM sodium fluoride and 0.5% (w/v) SDS and several passages through 26G ½ needles. Homogenates were then centrifuged 5 min at 13000 × g and 4°C to obtain the supernatants. Protein concentration was measured with the BCA kit.

### Electrophoresis and Western blot

HSA, serum or cellular extracts modified with AX or AX-treated with reducing or oxidizing agents and the corresponding unmodified controls were incubated in Laemmli buffer for 5 min at 95°C. Protein samples (1-15 μg) were loaded on SDS-PAGE gels and electrotransferred to Immobilon-P Transfer membranes for immunoblotting. Membranes were stained with Ponceau S (Sigma) as a control of protein loading and transfer. After destaining, membranes were incubated with 1:200 (v/v) mouse monoclonal anti-AX AO3.2 [16] and 1:2000 (v/v) anti-mouse IgG-HRP (Dako Cytomation) diluted in 1%(w/v) BSA in TTBS. Detection was carried out by chemiluminescence using the ECL system and the signal quantified utilizing image scanning and Scion Image software.

### Determination of glutathione concentrations

RPMI 8866 cells (10^6^) were treated as above. Cells were harvested by centrifugation and washed three times with PBS. The pellets were kept frozen at −80°C until use. The levels of glutathione reduced (GSH) and oxidized (GSSG) forms were measured using the enzymatic method of Tietze as described by Rahman et al. [17]. The extracts (250 μl) were diluted 1:10 (v/v) or 1:2 (v/v) for detection of GSH or GSSG, respectively.

### Statistical analysis

GraphPad Prism v. 5.0 (GraphPad Software) was used for statistical analysis of the data. Data are shown as average values from at least three experiments ± SEM. Experiments containing two groups of data were analyzed using Student’s *t*-test for independent samples; differences were considered significant when p < 0.05.

## Results

### Effect of oxidants or reducing agents on AX-protein adduct formation

Oxidative stress can act as a concomitant factor or as a mediator of drug adverse effects and influence drug-protein adduct formation [8, 14]. We have previously shown that AX forms adducts with HSA and other serum proteins [8]. Therefore, we were interested in the effect of oxidants and reducing agents on protein adduct formation by AX. Preincubation of purified HSA with DTT prior to AX addition led to a reduction in the levels of AX-HSA adducts, as detected by immunoblot with the anti-AX antibody (Fig. 1B). In contrast, preincubation with GSH did not significantly affect adduct formation, whereas incubation with GSSG increased adduct levels. When the assay was carried out with total serum, DTT tended to reduce, whereas GSH and GSSG tended to increase, AX-HSA adduct formation, although these effects were not statistically significant (Fig. 1C). These observations suggested that formation of AX-protein adducts in vitro can be modulated by several redox-active agents. Therefore, we set out to study these effects in more detail.

### Thiol-containing compounds induce a cyclization of AX with the formation of AX-diketopiperazine

Although some of the effects observed in Fig. 1 could be due to the influence of redox agents on the proteins themselves (see below), we also considered the possibility that these compounds might have an effect on the AX molecule. Therefore, the integrity of AX (1 equivalent) in the presence of DTT (2 equivalents) was investigated after overnight incubation in aqueous media using several characterization techniques (Fig. 2). The ATR-FTIR spectra of the mixture lacked the typical high infrared frequency β-lactam carbonyl band (C=O stretching >1780 cm^-1^) (Fig. 2A), which indicated cleavage of the β-lactam ring. ESI-MS analysis of the resulting mixture showed the same molecular ion than AX, but different fragmentations, providing evidence of an intramolecular rearrangement (Fig. 2B). The ^1^H-NMR spectra showed the disappearance of the doublet signals corresponding to the β-lactam ring protons (H5 and H6) of AX at 5.39 and 5.35 ppm (J^5,6^ = 3.9 Hz), and their appearance at 4.02 and 5.17 ppm (J^5,6^ = 2.6 Hz) (Fig. 2C). The new positioning of these bands is in agreement with the formation of the DKP cyclic product in quantitative yields. In addition, the presence of oxidized and reduced DTT was detected. Moreover, the singlets corresponding to methyl groups of the C2 thiazolidine shifted from 1.41 and 1.35 ppm in AX to 1.52 and 1.16 ppm in DKP. This rearrangement was not observed when AX was incubated under the same conditions but without DTT. This observation, together with the fact that 1 equivalent of DTT is not consumed in the reaction, indicated that intramolecular amide formation by nucleophilic attack of the AX side chain amino group to its electrophilic β-lactam carbonyl occurs only in the presence of DTT; and that the reducing agent acts in a catalytic manner, through the formation of an intermediate thioester, as depicted in the mechanism schematized in Fig. 3A (see below)

**Figure 2.**
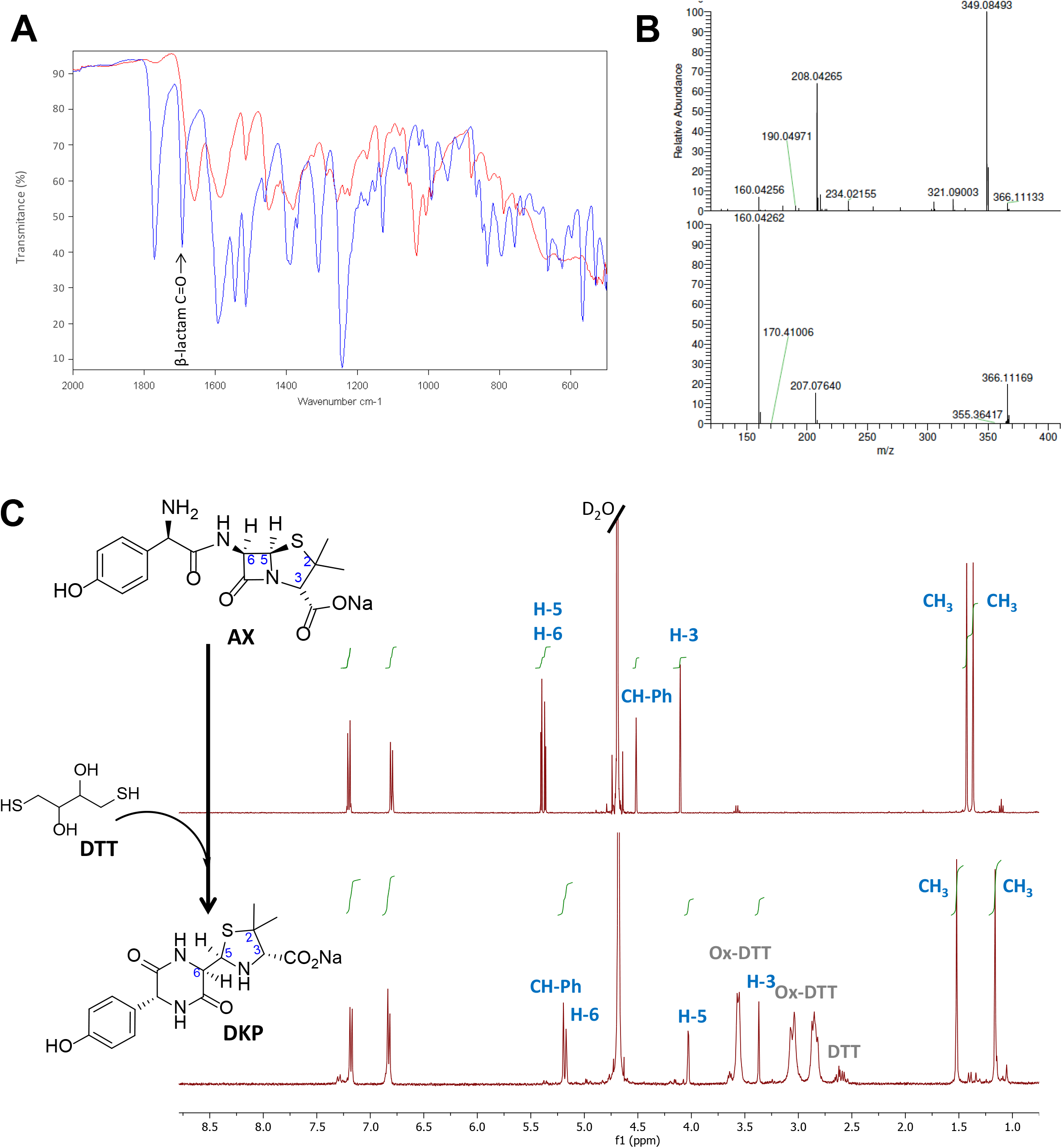
Monitoring amoxicillin reactivity in the presence of DTT in aqueous solution. **(A)** Attenuated total reflection-Fourier transform infrared (ATR-FTIR) spectra of the starting β-lactam, amoxicillin (AX, blue) and the resulting diketopiperazine (DKP, red) after DTT incubation. The band corresponding to C=O stretching of β-lactam (>1780 cm^-1^) in the AX molecule disappeared after reaction with DTT, which indicated its cleavage, in agreement with the formation of DKP. **(B)** Electrospray ionization-mass spectrometry (ESI-MS) of the starting β-lactam AX m/z (%): 366 (M+, 5), 349 (100), 208 (66), upper panel, and of the resulting DKP m/z (%): 366 (M+, 22), 207 (16), 160 (100), showing the same molecular ion peak than AX, but different fragmentation pattern and base peak, lower panel. **(C)** ^1^H-NMR spectrum of the starting β-lactam AX (in D_2_O), upper panel, and of the resulting DKP obtained after incubation of DTT (2 equivalents) with AX (1 equivalent) in H_2_O overnight at room temperature, lower panel. The resonances of β-lactam protons (H5 and H6) were used to monitor the reaction, shifting from 5.39 and 5.35 ppm to 4.02 and 5.17 ppm. The DKP spectrum also shows peaks corresponding to reduced and oxidized forms of DTT in the 2.5 to 3.7 ppm range.

To investigate whether this reactivity takes place under conditions closer to those of the physiological environment, the incubation experiments were carried out in PBS buffer at 37°C using different AX:DTT ratios (Table 1), and the progress of the reaction was monitored by ^1^H-NMR using the chemical shifts for AX β-lactam ring protons (H5 and H6). In all cases, we observed a time-dependent reduction in AX signals together with the appearance of signals consistent with the formation of the DKP cyclic product (H5 and H6, at 4.12 and 5.26 ppm, respectively) (Fig. S1). The amoxicilloyl thioesther intermediate might undergo an intramolecular aminolysis favored by the formation of the six-membered heterocycle as soon as it is formed. Moreover, DKP formation was shown to be time- and DTT concentration-dependent. We also considered the possibility that the reaction, instead of being catalyzed by the amoxicilloyl thioesther formation, could be favored by the redox properties of DTT. With the aim of investigating this potential mechanism, the fate of AX incubated in the presence of ethanothiol, as the nucleophile, was studied. This treatment resulted in virtually complete transformation of AX into DKP (Fig. S2), which confirmed the thiol-catalyzed intramolecular aminolysis of the AX β-lactam. Of note, as the DKP produced possesses the same mass than AX, the cyclization of the antibiotic in the presence of thiol-containing compounds would remain undetected if monitored only by MS [18].

**Table 1.**
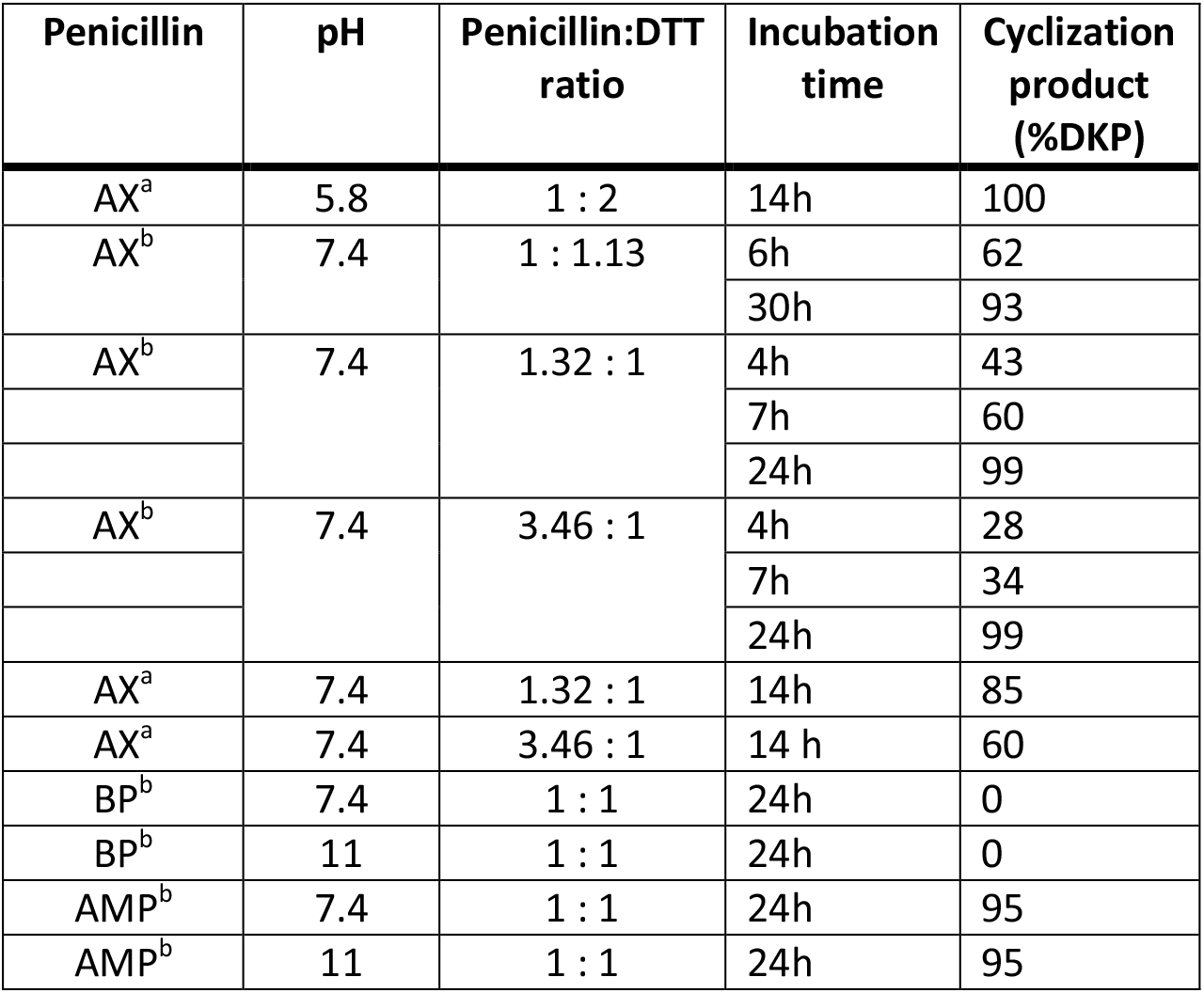
Reactivity of penicillins in presence of DTT in different conditions. Incubations were carried out at 37°C for the time stated in each case in (a) distilled water or PBS and then lyophilized; or (b) deuterated solvent (D_2_O, PBS or carbonate). In both cases, ^1^H-NMR was employed for identifying and quantifying the resulting cyclization product.

Based on these data, we propose the cyclization mechanism represented in Fig. 3A, in which the formation of DKP would be dependent on the presence of the amino group in the lateral chain of AX. To confirm this, the reactivity of other penicillins (with identical nuclear bicycle structure, and similar or different side chains) and cephalosporins (with different nuclear bicycle structure and identical, similar or different side chains) was studied in the presence of DTT (2 equiv) at neutral or basic pH. Among aminopenicillins, AMP was chosen since it shares the nuclear bicycle structure of AX and the presence of amino functionality in its side chain. AMP incubation with DTT at neutral pH for 24 h resulted in the formation of its DKP-AMP, showing the same reactivity than AX through intramolecular cyclization (Figs. 3A and S3). Next, BP was used as a model penicillin with the same nuclear bicycle structure, but without amino side chain functionality. In contrast to AX and AMP, the spectrum of BP suffered no modification after incubation with DTT at neutral pH (Fig. S4). However, addition of DTT at basic pH clearly elicited BP hydrolysis to the benzylpenicilloic acid and the appearance of small amounts of the thioesther intermediate. Thus, the mechanism of the reaction in this case would correspond to that depicted in Fig. 3B. In contrast, no reactivity was observed upon addition of DTT to cephalosporins bearing identical side chains than AX (cephadroxil) or AMP (cefaclor) at neutral or basic pH (Figs. 4, S5 and S6). Additionally, incubation of cefuroxime, a cephalosporin without free amino functionality in the side chain, in the presence of DTT at physiological pH resulted in only a small increase of the rate of hydrolysis compared to the incubation without the thiol compound. Moreover, the extent of the hydrolysis did increase at basic pH when using DTT (Figs. 4 and S7).

**Figure 3.**
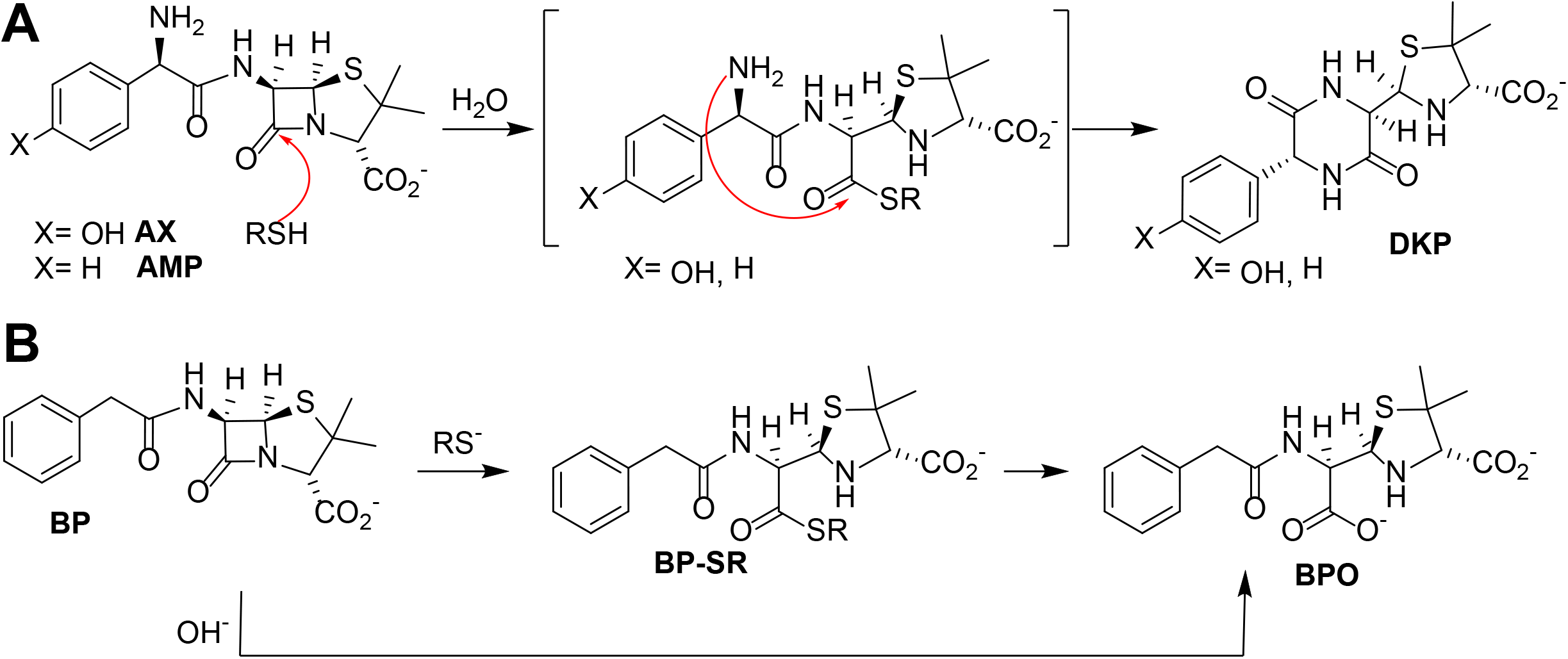
Reactivity of different penicillins in the presence of thiols. **(A)** Nucleophilic ring opening of aminopenicillins – amoxicillin (AX) and ampicillin (AMP)–by thiols and subsequent intramolecular aminolysis resulting in their corresponding diketopiperazine (DKP). **(B)** Hydrolysis of benzylpenicillin (BP) catalyzed by thiols, with formation of the benzylpenicilloate thioester intermediate (BP-SR), yielding the benzylpenicilloate (BPO).

**Figure 4.**
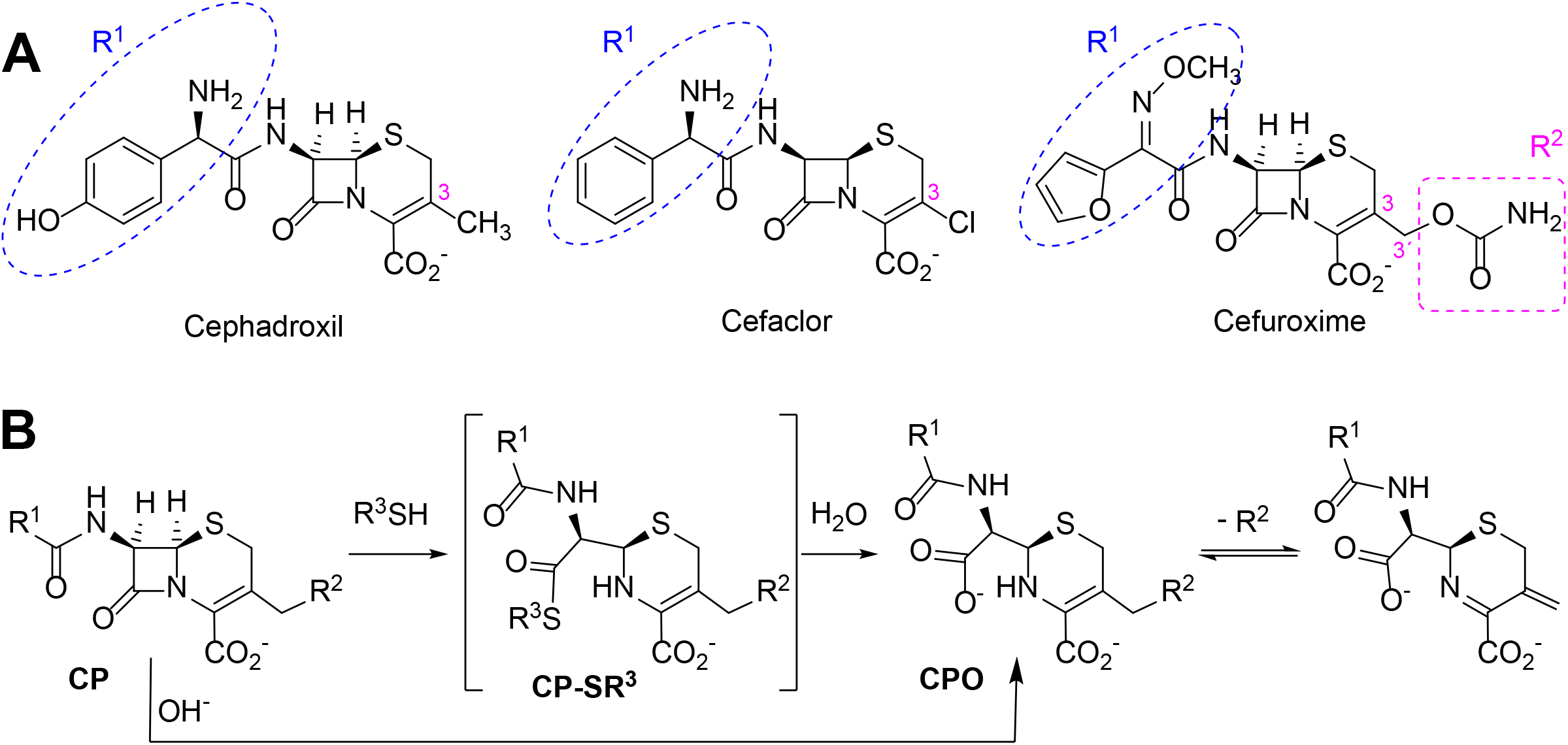
Structure and reactivity of cephalosporins in the presence of thiols. **(A)** Structure of different cephalosporins included in the study with highlighted R^1^ and R^2^ substituents. **(B)** Scheme of proposed reactivity of cephalosporins (CP) with a good R^2^ leaving group in presence of thiols, resulting in the thioester intermediate (CP-SR^3^) and the subsequent cephalosporoate (CPO), whose expulsion of the leaving group at C3’ affords the exo-methylene compound.

### Interaction of biologically or pharmacologically relevant thiol-containing molecules with AX

In view of the data obtained with DTT, we explored whether other relevant thiol-containing compounds could induce AX transformation (Table 2). The antioxidant N-acetyl-L-cysteine, which is the active principle of widely used mucolytic pharmacological preparations, also promoted DKP formation. Importantly, GSH, the main low molecular weight intracellular thiol antioxidant, effectively induced cyclization of AX. In contrast, nitro-benzyl-glutathione, a GSH analog in which the thiol group is protected, and GSSG, did not elicit this effect.

**Table 2.**
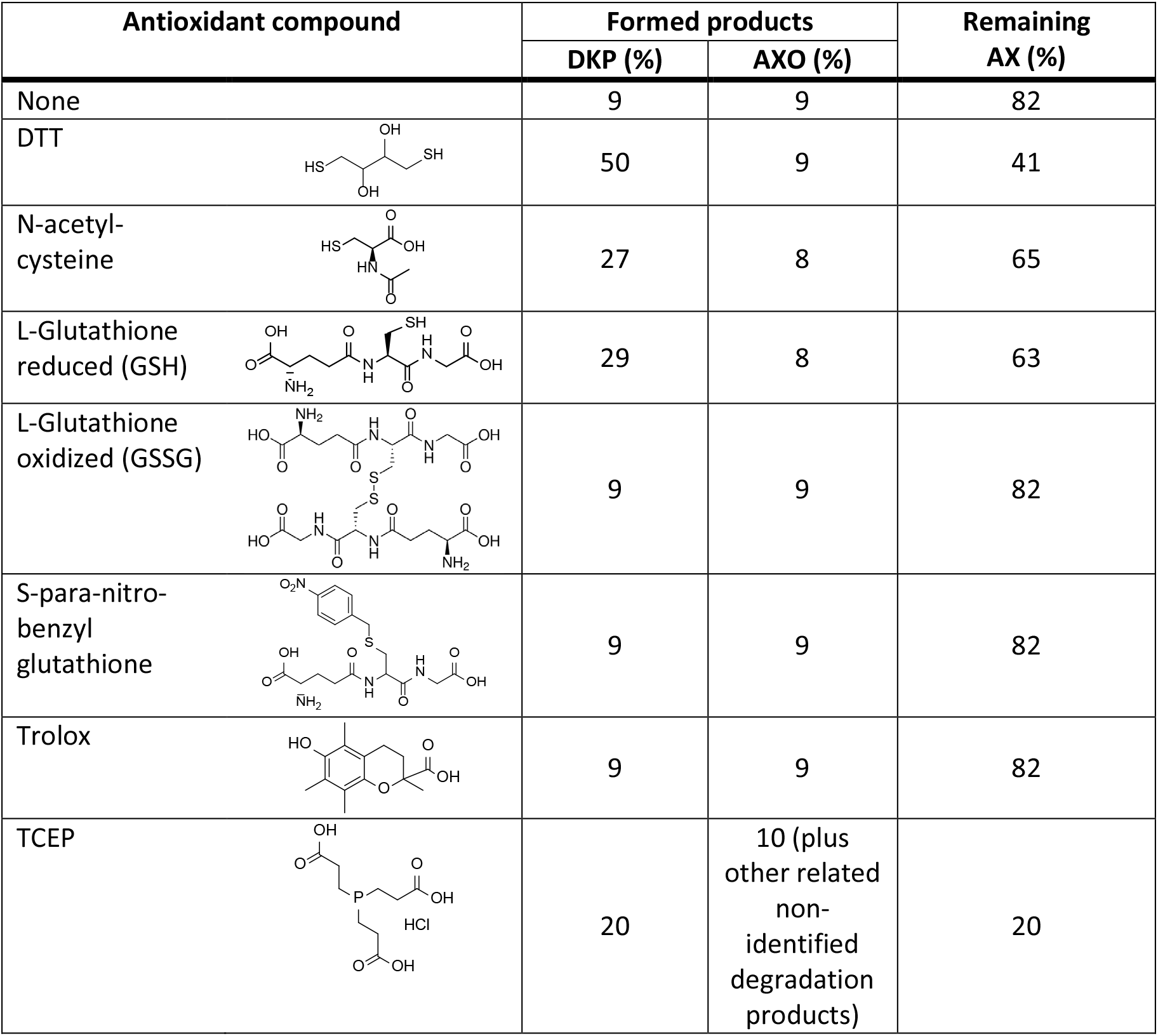
Reactivity of AX with thiol (protected, reduced or oxidized forms) and non-thiol antioxidants. Incubations were carried out using equimolar amounts of AX and the thiol functional-reagent or non-thiol antioxidant, in PBS at 37°C for 24 h and lyophilized. Integration of ^1^H-NMR signals was employed for calculating the quantity of obtained products.

| Antioxidant compound | Formed products |  | Remaining AX (%) |
| --- | --- | --- | --- |
|  | DKP (%) | AXO (%) |  |
| None | 9 | 9 | 82 |
| DTT | 50 | 9 | 41 |
| N-acetyl-cysteine | 27 | 8 | 65 |
| L-Glutathione reduced (GSH) | 29 | 8 | 63 |
| L-Glutathione oxidized (GSSG) | 9 | 9 | 82 |
| S-para-nitro-benzyl glutathione | 9 | 9 | 82 |
| Trolox | 9 | 9 | 82 |
| TCEP | 20 | 10 (plus other related non-identified degradation products) | 20 |

### Effect of non-thiol based antioxidants on AX availability

In view of the above results it was of interest to explore whether non-thiol based antioxidants commonly used in protein biochemistry and cell biology could interact with AX. This knowledge will be necessary for any subsequent studies involving the interaction of AX with proteins which require the presence of an antioxidant. The antioxidant Trolox did not alter AX (Table 2). In contrast, the widely-used antioxidant TCEP, induced decomposition of AX into several products, including DKP, amoxicilloate and other non-identified substances, which accounted for up to 80% of the resulting mixture. In addition, it should be taken into account that TCEP itself can suffer some decomposition in phosphate buffers at neutral pH [19]. Therefore, Trolox appears as a suitable non-thiol antioxidant for AX-protein studies.

### AX transformation into DKP implies a loss of reactivity and antibacterial potency

DKP is a known metabolite of AX, considered to have lower or residual reactivity and bacteriostatic activity [18], and which has been detected in humans, in experimental animals and in the environment [18, 20, 21]. To confirm the reduced DKP reactivity its interaction with butylamine, as a model of a nucleophilic compound susceptible to adduction by AX, was explored. ^1^H-NMR analysis of incubation mixtures containing equimolar amounts of DKP and butylamine revealed quantitative preservation of DKP (95% of the starting amount), thus confirming its lack of reactivity.

Next, the effect of preincubating AX with DTT or non-thiol antioxidants on its capacity to covalently modify HSA was monitored. For this purpose, antioxidant concentrations within the range used in cellular or biochemical applications were included in the assays. In particular, low micromolar to millimolar concentrations of Trolox are used in cell culture and in vivo applications [22], whereas low millimolar TCEP concentrations are mostly utilized in protein preparations [23]. DTT is generally included at low millimolar levels in most cell lysis and protein extraction protocols and at high millimolar concentrations in protein reduction methods. Therefore, freshly prepared mixtures of AX with vehicle (PBS) or the indicated antioxidants (0 h) or mixtures that had been preincubated for 24 h at 37°C (24 h) were combined with a solution of HSA and the formation of AX-HSA adducts was monitored 24 h later. As observed in Fig. 5A, AX preincubation with PBS or Trolox did not significantly reduce AX-HSA adduct formation with respect to the adducts formed by the freshly prepared mixtures, whereas preincubation with TCEP showed a tendency towards decreasing adduct levels. This behavior is consistent with the lack of interaction of Trolox with AX and the complex interaction exhibited by TCEP with the antibiotic as revealed by ^1^H-NMR (Table 2). Remarkably, the formation of adducts by the freshly prepared DTT-AX mixture was clearly lower than adducts formed by AX in PBS (control). Moreover preincubation of AX with DTT further diminished the capacity of AX to form adducts with HSA (Fig 5A).

**Fig. 5.**
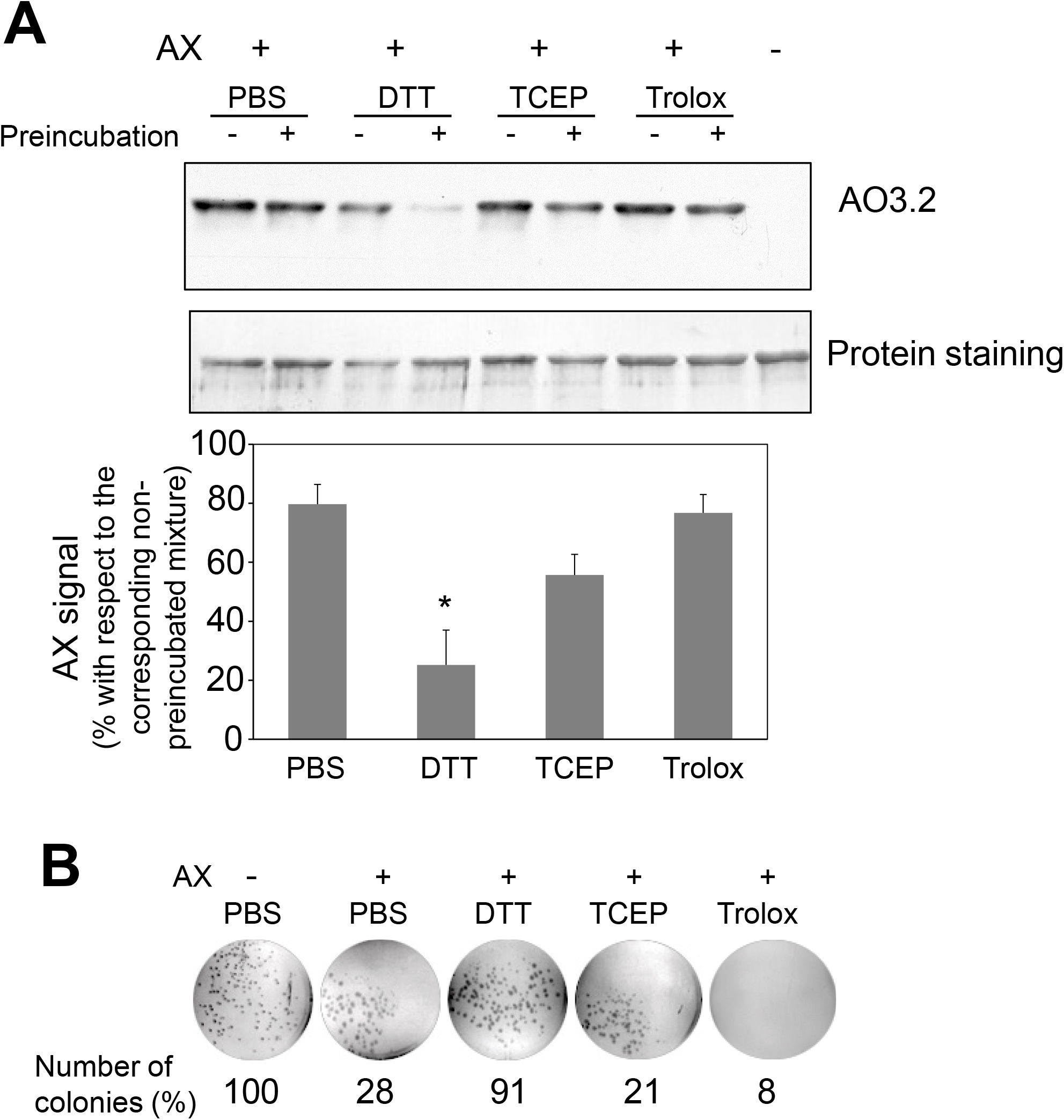
Effect of various antioxidants on the capacity of amoxicillin to form protein adducts and inhibit bacterial colony formation. **(A)** Mixtures of amoxicillin (AX) at 5 mg/ml (14 mM) and 100 mM DTT, 1 mM TCEP or 1 mM Trolox in PBS were either prepared fresh or preincubated for 24 h at 37°C. All mixtures were combined with an equal volume of HSA at 0.6 mg/ml in PBS and subsequently incubated for 24 h at 37°C. Finally, aliquots containing 4 μg of HSA were separated by SDS-PAGE and the formation of AX-HSA adducts was detected by western blot using anti-amoxicillin AO3.2. The blot shown corresponds to a representative experiment out of three with similar results. The histogram depicts the mean ± SEM of the intensity of the AX-HSA adduct signal obtained with every preincubated mixture with respect to that obtained with the freshly prepared mixture (100%) for each of the agents tested.(*p≤0.05). **(B)** AX was incubated in the presence of vehicle or reducing agents as above, and incubation mixtures were used to prepare LB-agar plates so that the theoretical final concentration of AX would be 100 μg/ml. *E. coli* DH5α were plated and allowed to grow for 24 h at 37°C, after which, colonies were imaged on a transilluminator and counted. Results show the number of colonies observed as percentage of those grown in the absence of AX, and are from a typical experiment out of three carried out.

Interaction of AX with thiols also reduced its antibacterial potency, as evidenced using two different approaches, namely, evaluation of colony formation and of the growth of suspension cultures of AX-sensitive *E. coli*. Preincubation of AX with PBS or the non-thiol antioxidants TCEP and Trolox did not impair its antibacterial effect since strong inhibition of colony formation was observed (Fig. 5B). In contrast, incubation with DTT relieved this inhibition, thus indicating its interference with the bacteriostatic action of AX (Fig. 5B). Additionally, assessment of the effect of thiol-containing compounds on the antibacterial action of AX by following *E. coli* growth in turbidity assays showed that all the thiol-containing compounds, namely, DTT, GSH and 2-mercaptoethanol, attenuated the bacteriostatic effect of AX in a concentration-dependent manner (Fig. 6). DTT was the most potent, blocking AX effect at equimolar ratios (Fig. 6A), as expected from its higher reduction potential and the presence of two thiols in the molecule. On the other hand, GSH and 2-mercaptoethanol displayed similar potency, needing thiol:AX ratios above 4:1 to obtain a complete abolition of antibiotic effects (Fig. 6B and C).

**Figure 6:**
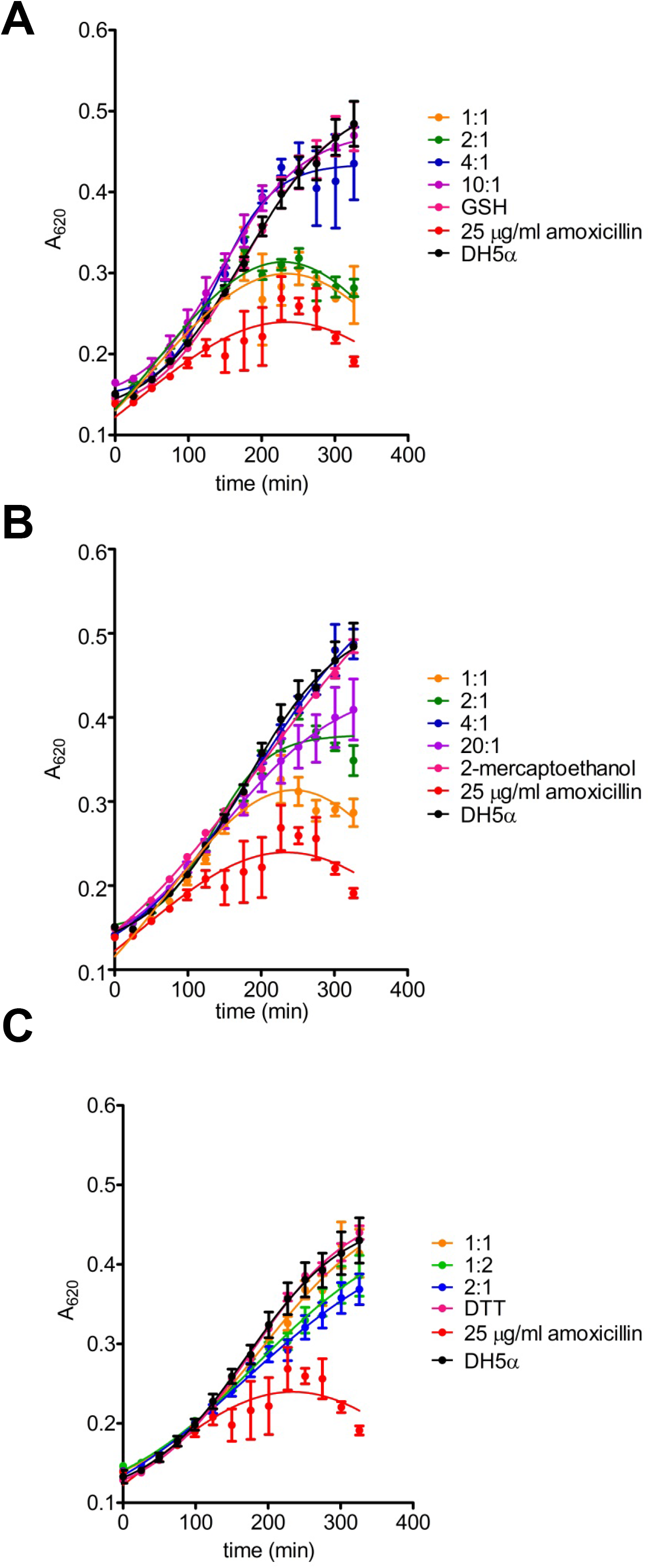
Effects of amoxicillin inactivation by reducing agents on bacterial growth. The ability of glutathione **(A)**, 2-mercaptoethanol **(B)** and DTT **(C)** to inactivate amoxicillin (AX) was determined after incubation for 24 hours at 37°C with the reducing agent vs. AX molar ratios indicated in the figure. The turbidity assay was carried out on a multiwell plate using *E. coli* DH5α and a final theoretical AX concentration of 25 μg/ml per well. Controls of bacterial growth without the antibiotic, with 25 μg/ml AX or appropriate concentrations of each reducing agent were also included; only controls with the highest concentrations of reducing agents are shown for clarity. The figure depicts results of a typical experiment (n=3) carried out in triplicate (mean ± SEM, *p ≤ 0.05).

### Importance of AX-thiol interaction in biological settings

The interactions between AX and thiols could be highly relevant for the action of this antibiotic in biological environments, in which levels of thiol-containing compounds have a very important role in the control of the redox status. Among them, GSH reaches millimolar concentrations in cells. GSH synthesis can be blocked by BSO, an inhibitor of γ-glutamylcysteine synthetase [24], the rate limiting enzyme of this pathway, thus reducing cellular GSH levels and inducing oxidative stress. In order to explore the interplay between redox status and AX action, we treated RPMI 8866 cells with AX alone or in combination with BSO and the effect on GSH concentrations was measured. AX alone did not alter cellular glutathione levels, neither the GSH/GSSG ratios (Fig 7A). In contrast, when cells had been pretreated with BSO before AX addition, a drastic decrease in both GSH and the GSH/GSSG ratio was observed compared to both control and AX-treated cells (Fig 7A). As mentioned before, AX can form adducts with cellular proteins that can be detected by immunological methods [9, 10]. Interestingly, reduction of GSH levels and the GSH/GSSG ratio in RPMI 8866 cells by pretreatment with BSO correlated with a significant increase in the formation of AX-protein adducts (Fig 7B). Nevertheless, whether this effect is related to an improved availability of AX due to decreased GSH levels requires further study.

**Figure 7.**
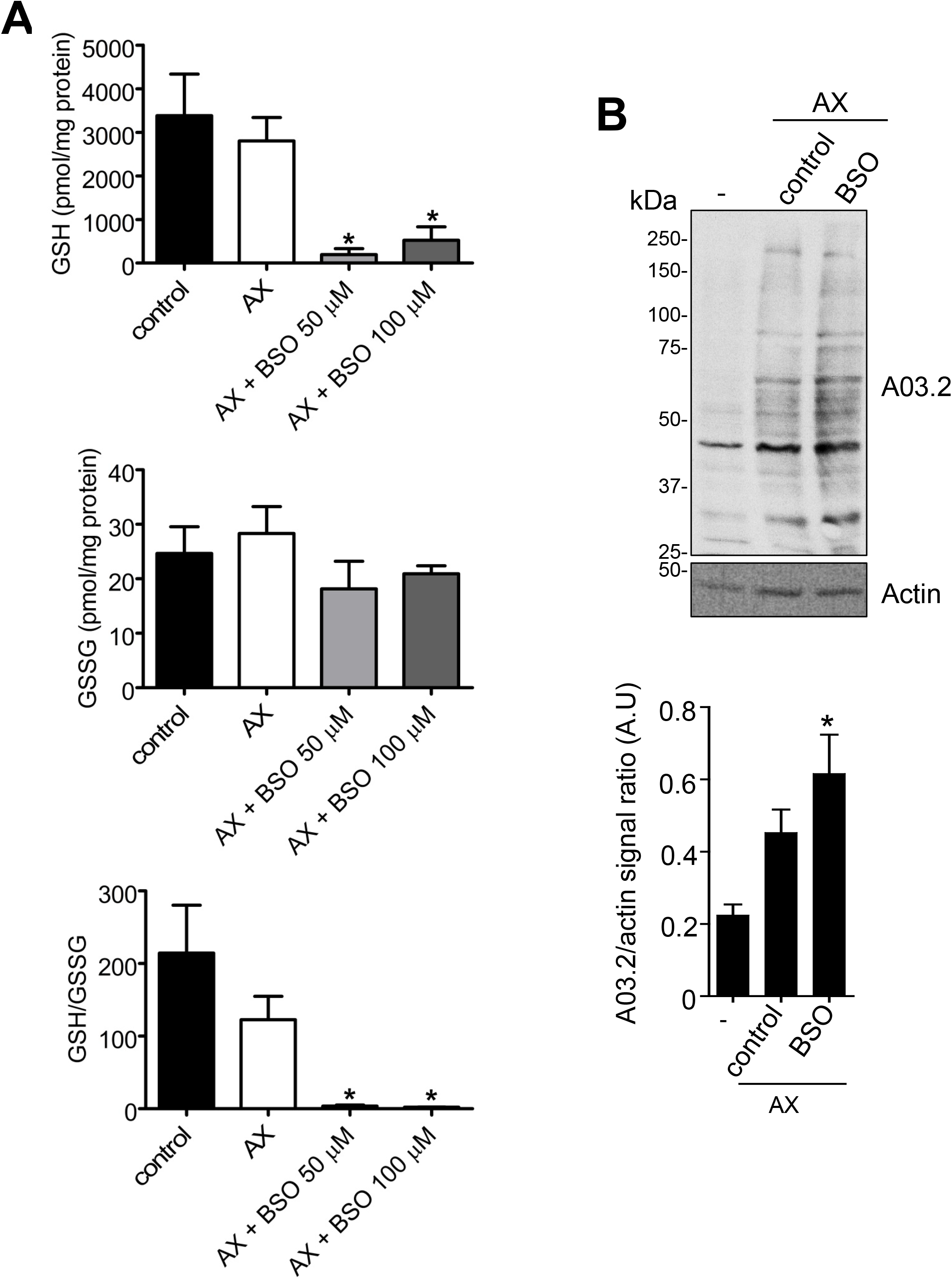
Effects of amoxicillin and glutathione availability in a cellular context. **(A)** RPMI 8866 cells (10^6^) were pre-treated with PBS (control), or buthionine sulfoximine (BSO 50 or 100 μM) for 24 h, after which, either PBS or amoxicillin (AX), as indicated, were added for 16 h, and the intracellular concentrations of reduced (GSH) and oxidized glutathione (GSSG) at the end of the incubation measured. The figure shows results (mean ± SEM) of five independent experiments that were measured in triplicate. The resulting GSH/GSSG ratio is also depicted. **(B)** Impact of inhibiting GSH synthesis in RPMI 8866 cells with 50 μM BSO on the formation of AX-protein adducts. Cells were treated with or without BSO and AX as above, harvested and the incorporation of AX into proteins analyzed by western blot using anti-amoxicillin AO3.2. The AX signal of each lane was quantified by densitometric scanning and corrected using actin levels as loading control. The graph shows the mean ± SEM of seven independent experiments (*p<0.05 vs control AX).

## Discussion

AX is a widely used antibiotic and one of the most frequently involved in allergic reactions. Therefore, AX reactivity and conjugation to carrier molecules have been the subject of thorough investigation [15, 25]. The redox status is a major determinant of drug action and oxidative stress is a well-recognized coadjuvant factor in drug adverse reactions [14]. Moreover, the redox status and oxidative modifications can influence drug binding to carrier proteins. Control of redox status is brought about by a complex interplay of enzymatic mechanisms and small molecule antioxidants. The tripeptide GSH, which bears a strong nucleophilic thiol group, plays a key role in this regulation in cells, and it also participates in drug detoxification [26]. Besides GSH, drugs can encounter a variety of thiol-containing molecules along their administration or distribution route in the body, including peptides, proteins or other drugs, like mucolytic agents containing N-acetyl-L-cysteine. Here we report the interaction of thiol-containing molecules with AX. Our results constitute the first evidence for the catalytic inactivation of this antibiotic, and potentially other β-lactams of similar characteristics, by thiols. This finding could have both pharmacological and biotechnological applications.

From the chemical point of view, our data show that, in the presence of thiols, AX is transformed into its DKP. This indicates an intramolecular amidation subsequent to thiolysis (Fig. 3A). On the one hand, the chemical reactivity of penicillins is related to the electrophilic properties of the β-lactam carbonyl, which is associated with the high tension within the β-lactam ring fused to the thiazolidine ring [27]. Additionally, the protonated form of the thiol group is not particularly reactive, whereas the thiolate anion is nucleophilic and therefore can participate in reactions with electrophilic reactive species [28]. In this context, interaction between AX and thiol molecules may stablish an acid-base equilibrium, which generates catalytic amounts of thiolate anion. This initiates the reaction with the β-lactam ring towards the formation of an intermediate thioester, the geometry of which should favor further intramolecular amidation/cyclization (by the side chain amino group) [29]. To the best of our knowledge, the thiol catalysis for intramolecular reactivity forming cyclizated structures characterized here has not been previously reported in β-lactam antibiotics.

The existence of interactions between thiols and β-lactam antibiotics, both in vitro and in vivo, has long been known. Early studies detected a loss of the antibacterial activity of penicillin induced by cysteine, which was attributed to chemical reactions involving the thiol and/or amino groups of the amino acid [30–32]. More recent studies have described in detail the thiol-catalyzed hydrolysis of BP [33, 34], in which the catalytically reactive form of the thiol is the thiolate anion [33]. This result is in agreement with our observations, which indicate a higher reactivity between BP and the thiol at basic pH. The penicillin thioester, previously identified by ^1^H-NMR, is the first detectable product in the reaction, which subsequently undergoes hydrolysis to the benzylpenicilloic acid (Fig. 3B) [33]. Therefore, the opening of the strained four-membered ring is faster through thiolysis than hydrolysis due to the higher nucleophilicity of the thiolate anion in comparison with the oxygen anion. Moreover, the minute amount of the thioester intermediate detected supports its low stability and high reactivity towards hydroxyl anions. [33].

The six-membered ring to which the β-lactam is fused in cephalosporins, results in lower tension within the β-lactam ring compared to those of penicillins, and therefore less reactivity against thiolate anions [27]. Thiol-catalyzed hydrolysis of cephalosporins has been described for a series of these antibiotics with kinetics that depend on the basicity of the thiol [35]. Our observations indicate that the chosen aminocephalosporins, cephadroxil and cefaclor, are not reactive towards thiols at either neutral or basic pH. Results obtained with cephadroxil corroborate a previous publication stating that the nature of the C3 substituent influences thiolysis. The R^2^ chemical structure can modulate this reactivity depending on its ability to polarize the electronic binding [27]. In cephadroxil, position 3 is occupied by a methyl group, whose non-polar nature can explain the lack of effect we observe on the kinetic hydrolysis of the β-lactam ring (Fig. 4, S5). Cefaclor has a chlorine atom at position 3, whose high electronegativity should facilitate the opening of the β-lactam ring by electronic induction. Although in basic media we observe a low degree of hydrolysis of cefaclor leading to unidentified degradation products, none of them can be attributed to the thiol reactivity (Fig. 4, S6). Cefuroxime shows reactivity of the β-lactam group in the presence of thiols, resulting in its eventual hydrolysis and further degradation (Fig. 4, S7). This increased β-lactam reactivity is explained by the R^2^ substituent that may act as a leaving group [27], which is in agreement with previous observations in cephalosporins with a good leaving group at this position [35].

In biological settings and biochemical research, low molecular weight thiols are important antioxidants, which act through a variety of mechanisms. Among them, their role as components of the general thiol/disulfide redox buffer, metal chelators, radical quenchers, substrates for specific redox reactions (GSH), and specific reductants of protein disulfide bonds (thioredoxin) can be highlighted [36]. Thiol groups need to be deprotonated to form charged thiolates with higher reactivity compared to thiols. Thus, the inherent reactivity of low molecular weight thiol compounds depends on the acid dissociation constant (pKa) of their thiol groups, i.e. the lower the pKa the higher the reactivity of the thiol [28]. Besides DTT, we have studied the ability of several biochemically or pharmacologically relevant antioxidants to catalyze AX inactivation. Both GSH and N-acetyl-L-cysteine bear thiol groups with similar pKa values (in the range of 9.2-9.6). Moreover, these pKa values are close to those of thiol groups in DTT (9.2 and 10.1). Hence, differences observed in vitro for AX inactivation with thiol-containing antioxidants can be ascribed to the different number of thiol groups in the molecules used, one in GSH and N-acetyl-L-cysteine and two in DTT. Consistent with these facts, our assays demonstrate a more efficient inactivation of AX by DTT than with either GSH or N-acetyl-cysteine. (Table 2)

In physiological settings, the nature and abundance of the thiol-containing molecules that AX will encounter depends on the environment (e.g. plasma or tissues), the cell type and even the nutritional status (e.g. cysteine content after a protein-rich ingestion). Among them, HSA is one of the main antioxidants in plasma, where its concentration reaches 4.6 g/dL [37]. HSA represents more than 50% of the antioxidant capacity of this fluid in normal subjects [38]. In addition to its role as scavenger of reactive species [39], the state of Cys34 of HSA is considered the major determinant of plasma redox status and represents about 80% of the free thiol groups in blood [40]. Importantly, the redox state of HSA is altered under various pathophysiological conditions, including diabetes, renal disease, and during exercise, in which the proportion of reduced HSA (mercaptalbumin, thiol group of Cys34 in its reduced form) versus oxidized HSA (cysteinylated), is decreased. Therefore, it could be speculated that the state of oxidation of HSA could influence AX availability. Nevertheless, the interactions between AX and HSA are far more complex, since HSA is a major target for haptenation by AX [8]. HSA is synthesized in the liver, where its production attains 10.5 g/day, and has a turnover time of ~25 days [37]. According to these data, only 8.5% of HSA is renewed daily, thus allowing a long exposure time to drugs, which can favor the formation of drug-HSA adducts, as those detected in vitro and in the serum of patients [8, 41], and facilitate the development of allergic reactions.

In contrast, plasmatic GSH concentrations are low (<4 μM) [42], whereas intracellular concentrations reach 1-5 mM in most cell types and 5-10 mM in hepatocytes [24, 43, 44]. Hence, a major role of GSH on AX availability could be envisioned in intracellular environments, where additional antioxidant systems could contribute to DKP production. In fact, the normal GSH/GSSG ratios of 100:1 will favor the reductive and conjugating activities of glutathione, whereas decreases in this ratio due to oxidative stress will impair these activities enhancing the availability of drugs and AX [45, 46]. Additionally, it could be hypothesized that cellular proteins with thiol-disulfide active sites, such as GSR, could contribute not only to reduce GSSG levels and to the normalization of the GSH/GSSG ratio, but also to decrease AX availability by a thiol-mediated mechanism. Importantly, the distribution of GSH and of protein antioxidant systems is not uniform inside cells. For instance, in HeLa cells, glutathione redox potentials are more reducing in the mitochondrial matrix than in the cytoplasm [45], whereas the activity of glutathione transferases can differ between cellular compartments, including cytosol, mitochondria and nucleus [47]. Therefore, some of these protein antioxidant systems, together will glutathione, may contribute thiols to reduce the effective concentration of AX within the cell and both, the system involved and the ratio of conversion may depend on the subcellular compartment where the reaction occurs. Notably, free radicals have been reported to destroy β-lactam antibiotics in aqueous solution [48]. Nevertheless, whether this can occur under pathophysiological conditions needs to be ascertained.

The interaction of thiol-containing drugs and β-lactam antibiotics has been also studied in vivo. Available evidence is conflicting and both potentiating and interfering effects have been described. Cysteine-containing compounds have been reported to reduce the potency of several antibiotics, including AX [49, 50]. In clinical settings, the presence of thiol-containing compounds such as N-acetyl-L-cysteine could interfere with the antibiotic itself diminishing its efficacy. Nevertheless, N-acetyl-L-cysteine could also potentiate antibiotics’ effects by favoring their access to the sites of infection due to its mucolytic action or its ability to interfere with biofilms [51, 52]. Therefore, the clinical benefit or interference of thiol compounds with the bacteriostatic effect of AX needs to be evaluated in every specific setting. Administration through different routes or in an asynchronous way, as recommended in practice, appears to be an advisable strategy.

In summary, our results indicate that thiols may alter AX availability, both in vitro and in cellular environments, thus constituting a major determinant for AX action under pathophysiological conditions. Additionally, the oxidative stress concurrent with many pathological situations could be an important contributor to the formation of the larger amounts of AX-protein adducts observed under this type of stress and that seem involved in AX adverse reactions. These finding can be relevant for understanding the mechanisms involved in hypersensitivity reactions to β-lactams.

## Supporting information

Pajares 2019 Supplemental material

## Acknowledgements

This work was supported by grant SAF2015-68590-R from MINECO/FEDER and RETIC Aradyal from ISCIII/FEDER RD16/0006/0021 to DPS; RD16/0006/0001 to MJT, RD16/0006/0024 to MB; grants CP15/00103 and PI17/01237 from ISCIII/ERDF and PI-0179-2014 from Andalusian Regional Ministry Health to MIM. AA holds a “Sara Borrell” research contract (CD17/0146) supported by ISCIII from MINECO (confounded by the European Social Fund (ESF)).

