## Supplementary material for "Amoxicillin inactivation by thiol-catalyzed cyclization reduces protein haptenation and antibacterial potency": Pajares 2019 Supplemental material

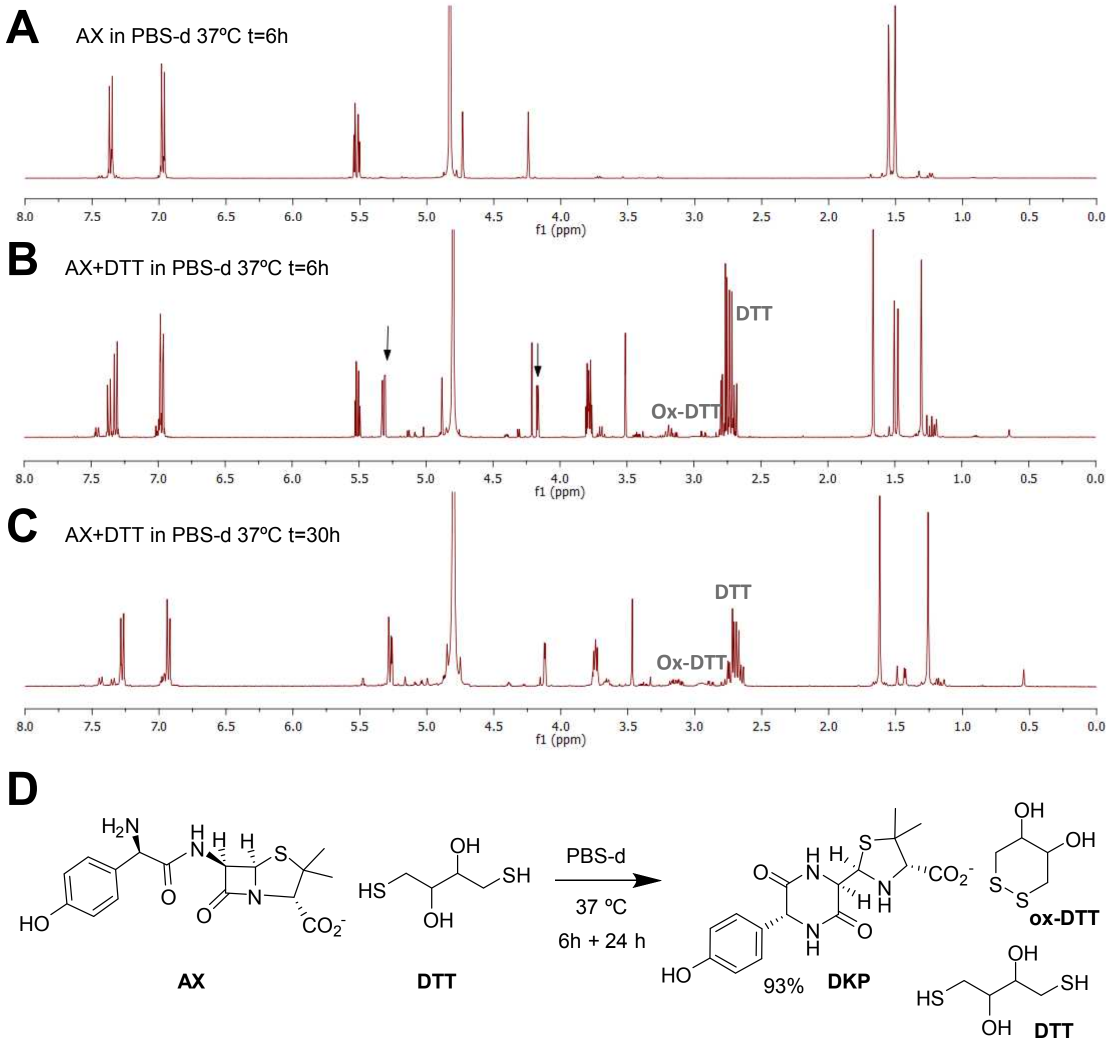


**Fig. S1. Monitoring amoxicillin reactivity in deuterated-PBS (PBS-d) at 37°C in the presence of DTT. (A)** Control reference of amoxicillin (AX) in PBS-d after 6 h, with no observed modification of AX; **(B)** AX (1 equivalent) incubation with DTT (1.13 equivalent) in PBS-d after 6h, with arrows pointing out H5 and H6 proton signals of the formed DKP structure. **(C)** AX (1 equivalent) incubation with DTT (1.13 eq) in PBS-d after 30 h, that shows only 7% of AX signals remaining and 93% conversion to DKP structure. 20% of the DTT is in oxidized form; **(D)** Reaction scheme of AX incubation with DTT.

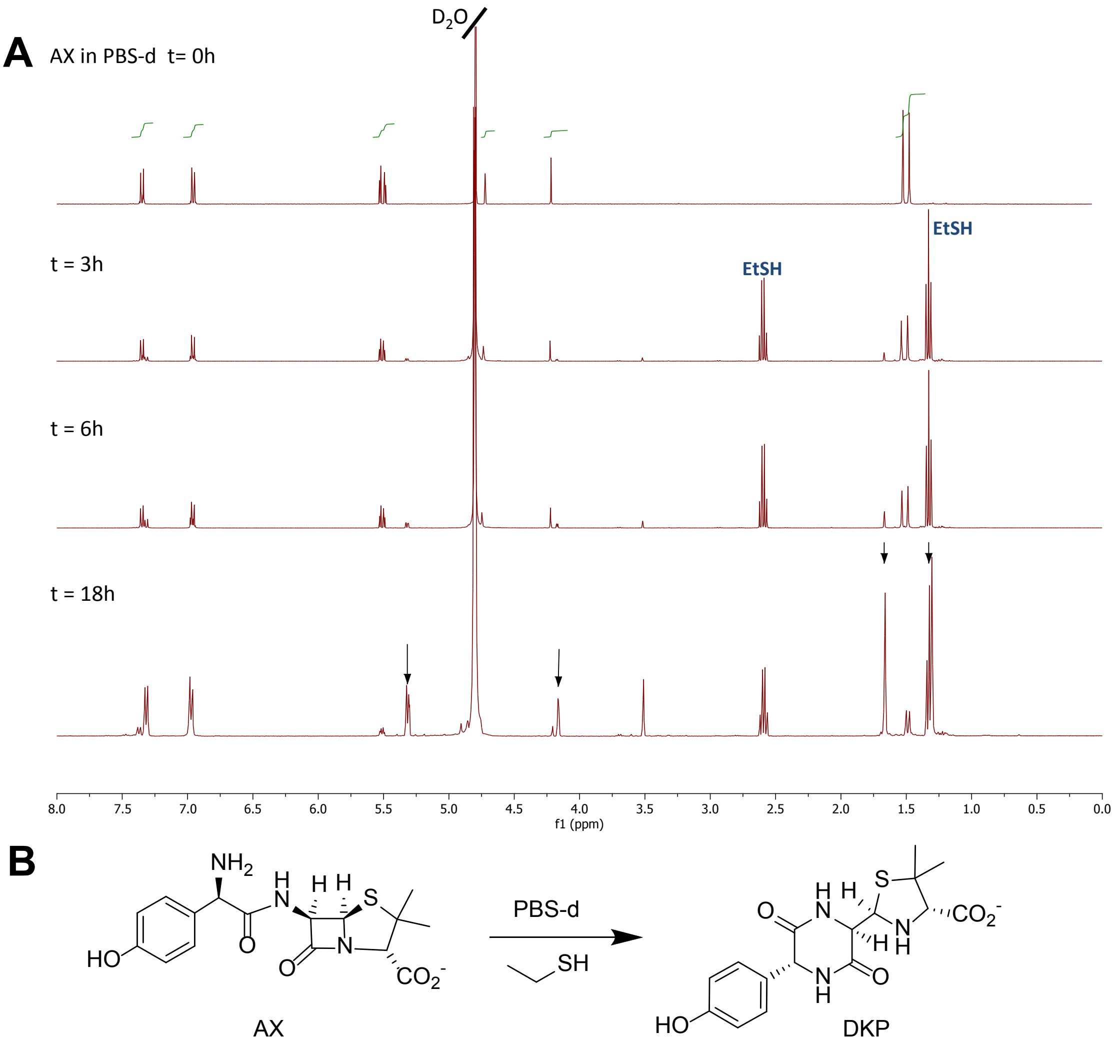


**Fig. S2. Monitoring AX integrity in the presence of an excess of ethanothiol (EtSH). (A)** The spectrum above shows the signals of the starting AX in PBS-d. The following spectra are registered after addition of an excess of EtSH to the AX solution in deuterated PBS (PBS-d) at 37°C at different times: 3, 6 and 18 h. Arrows point out H5 and H6 proton signals in the formed DKP structure and methyl groups of its thiazolidine ring. **(B)** Reaction scheme of AX incubation with EtSH in PBS.

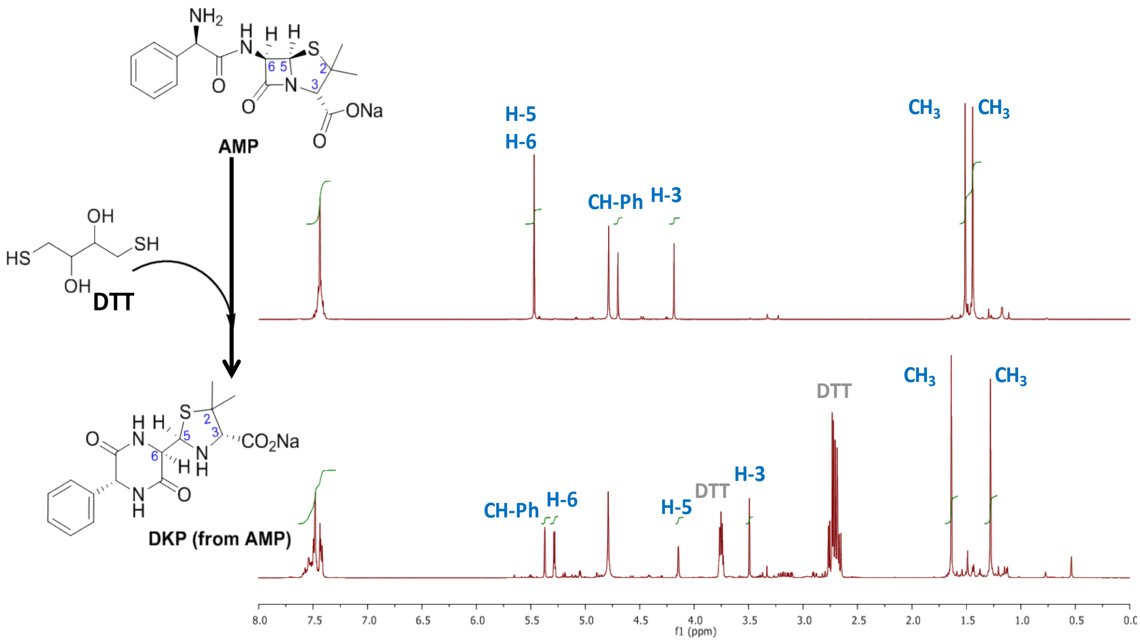


**Fig. S3. NMR monitoring of ampicillin (AMP) reactivity in the presence of DTT.** $^{1}H$ NMR spectra of AMP (above) and its resulting DKP (below) after incubation for 24h in the presence of 2 equivalents of DTT in $D_2O$. The resonance of β-lactam protons (H5 and H6) were used to monitor the reaction.

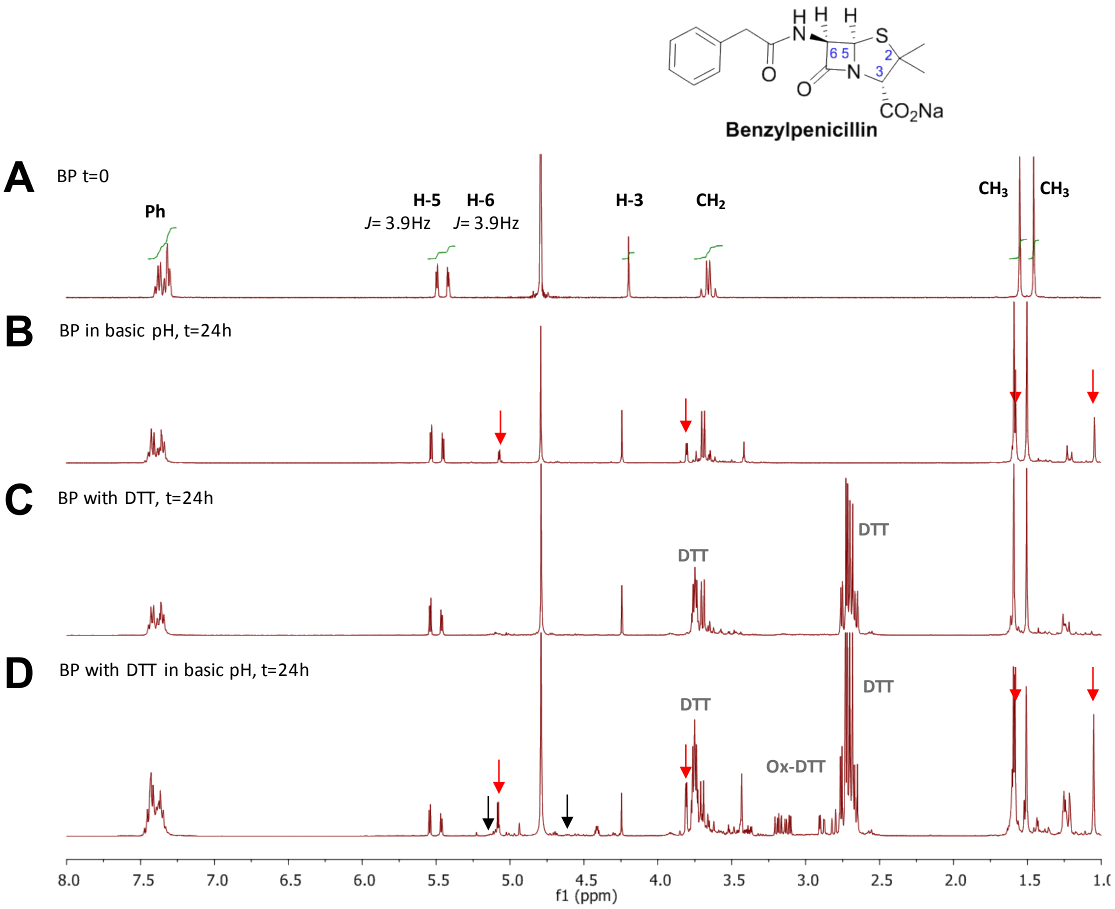


**Fig. S4. $^1$H-NMR monitoring of benzylpenicillin reactivity in the presence of DTT.** Spectrum of BP in $D_2O$ **(A)** at initial time and **(B-D)** at 24 h under different experimental conditions. **(B)** Spectrum of BP reactivity after 24 h in basic pH, which indicates the opening of the β-lactam. The resonance of β-lactam protons (H5 and H6) was used to monitor the progress of the reaction. Red arrows show the appearance of new chemical shift for H5 and H6 and methyl groups of benzylpenicilloate. Spectra **(C, D)** show the reactivity of BP with DTT at neutral and basic pH, respectively. The spectrum obtained in the presence of DTT at neutral pH **(C)** showed no modification of the starting BP. The spectrum obtained in presence of DTT at basic pH **(D)** showed signals corresponding to benzylpenicilloate, pointed out by red arrows, indicative of a high rate of hydrolysis in comparison with **(C),** due to the presence of the thiol reagent in basic media. Black arrows point out very small peaks that could correspond to the H5 and H6 protons of benzylpenicilloate thioester.

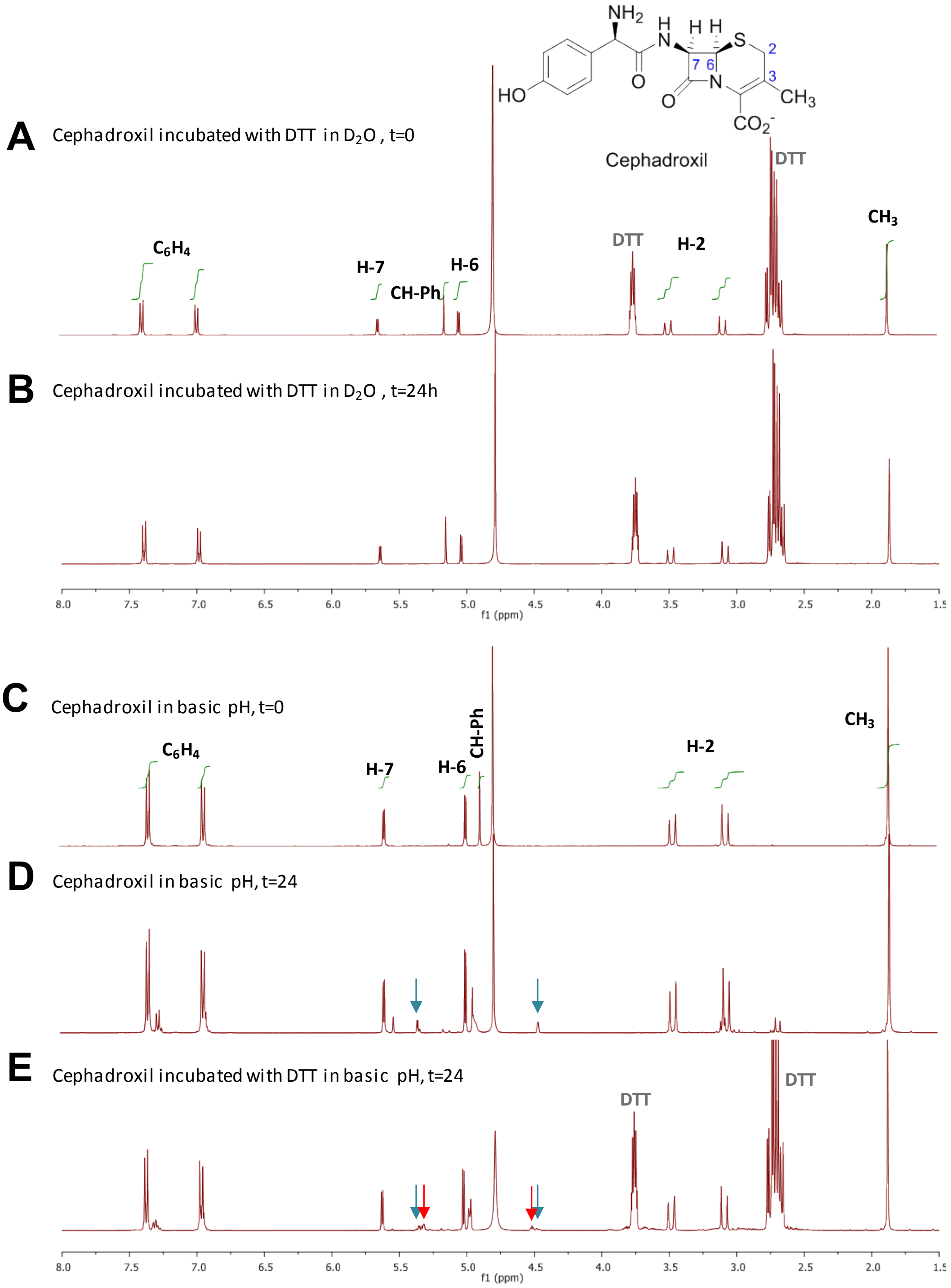


**Fig. S5. NMR monitoring of cephadroxil reactivity in the presence of DTT. (A)** $^1$H-NMR spectrum of the mixture of cephadroxil (1 equivalent) with DTT (2 equivalents) in $D_2O$ at initial time and **(B)** after 24 h of incubation. No modification was observed. **(C)** $^1$H-NMR spectrum of cephadroxil at basic pH and 0 h or **(D)** 24 h of incubation. Arrows show the appearance of small peaks corresponding to H7 and H6 of the cephalosporoate structure. **(E)** $^1$H-NMR spectrum of the incubation of cephadroxil (1 equivalent) with DTT (2 equivalents) for 24h in basic pH. Blue arrows show the appearance of peaks corresponding to the cephalosporoate (CPO) structure also formed in control experiment (**D**). Red arrows point out new small peaks that could be attributed to isomers of CPO or other degradation products.

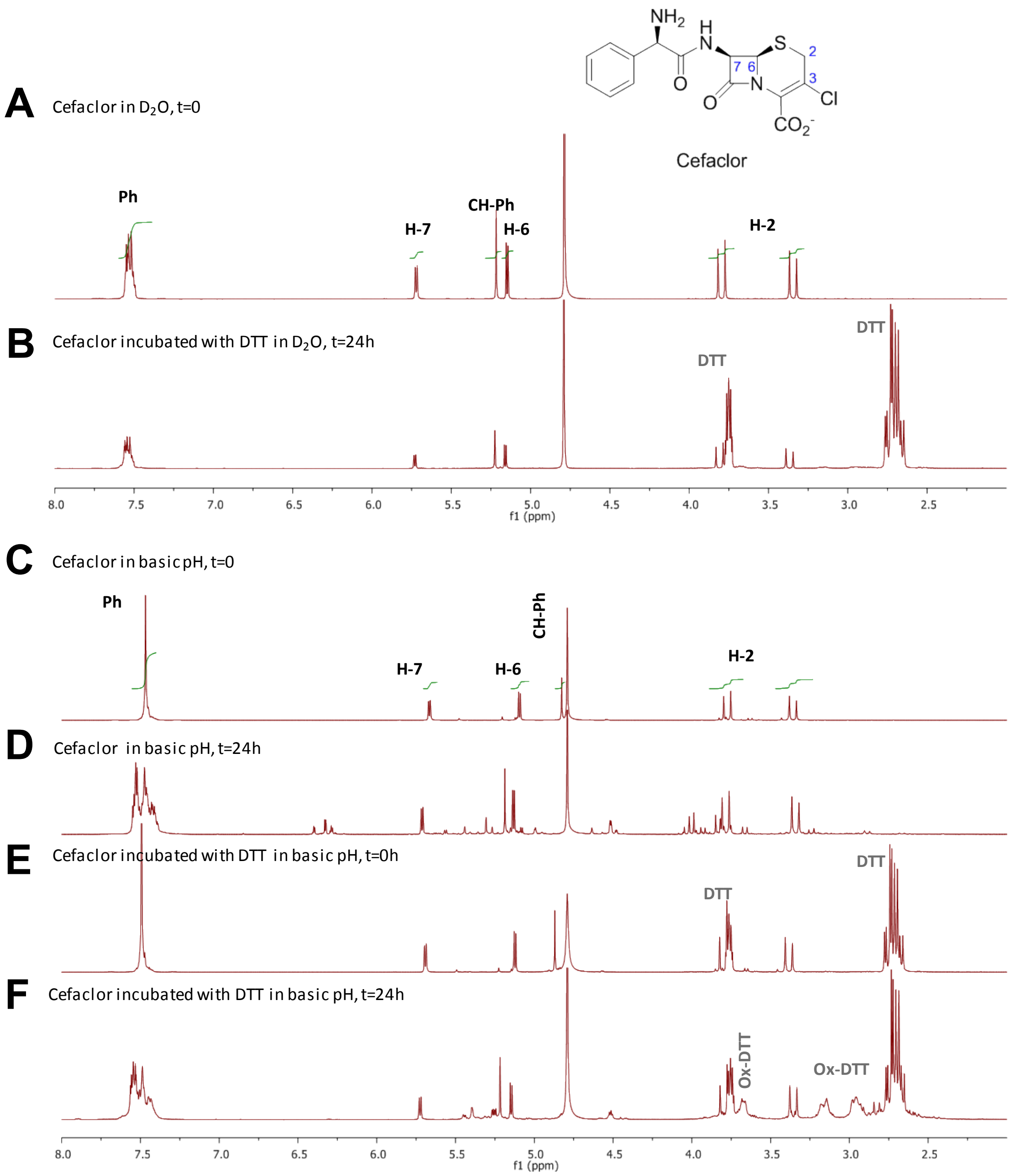


**Fig. S6. $^{1}$H-NMR monitoring of cefaclor reactivity in the presence of DTT. (A)** Spectrum of cefaclor in $D_2O$ at initial time. **(B)** Spectrum of the mixture of cefaclor (1 equivalent) with DTT (2 equivalents) in $D_2O$ after 24 h of incubation. No modification of the penicillin was observed. **(C)** Spectrum of cefaclor at basic pH at initial time. **(D)** Spectrum of cefaclor at basic pH after 24 h, where most signals correspond to intact cefaclor, although other peaks corresponding to degradation products are observed. **(E)** Spectrum of the mixture of cefaclor (1 equivalent) with DTT (2 equivalents) at basic pH at initial time. **(F)** Spectrum of the mixture of cefaclor (1 equivalent) with DTT (2 equivalents) at basic pH after 24 h of incubation, where most signals correspond to intact cefaclor, although other peaks corresponding to unidentified degradation products are observed, similar to those obtained with cefaclor only under basic conditions **(E)**. Oxidized and reduced forms of DTT are also observed.

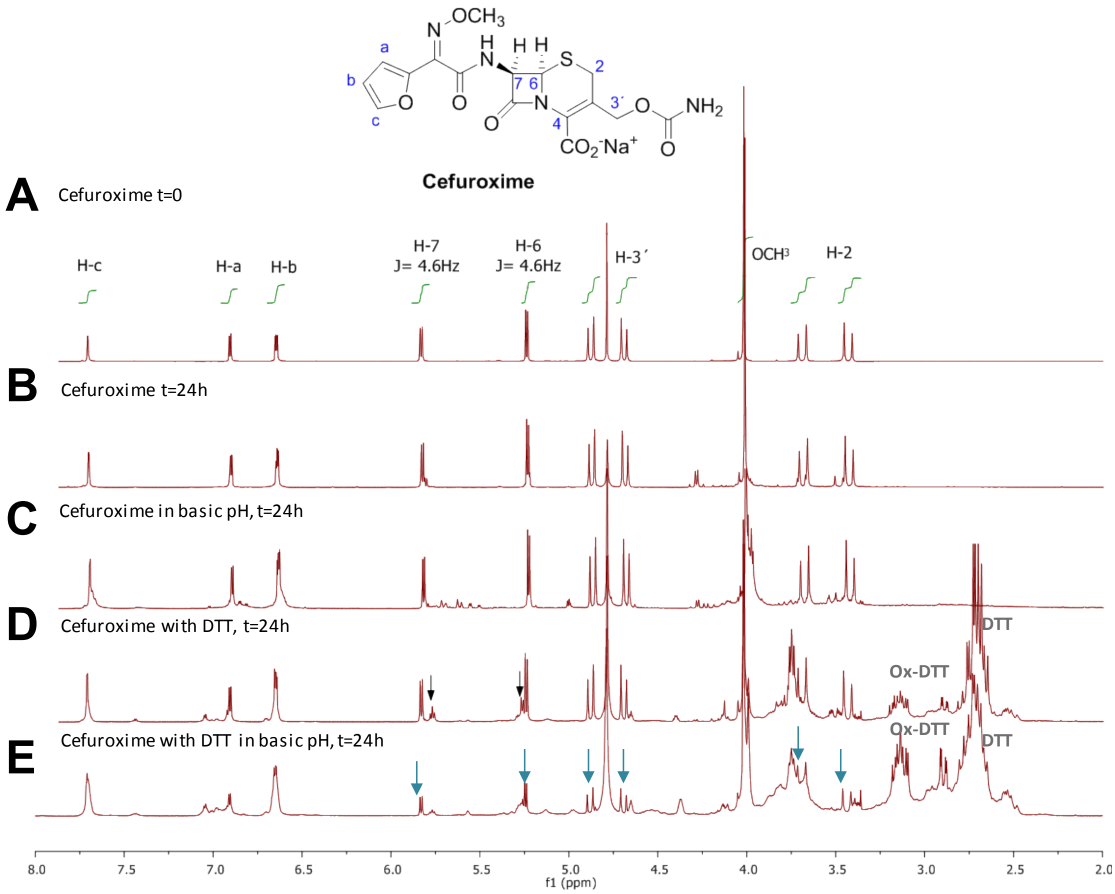


**Fig. S7. [1]H-NMR monitoring of cefuroxime reactivity in the presence of DTT. (A)** Spectra of cefuroxime in $D_2O$ at initial time and **(B-E)** after 24 h incubation under different experimental conditions. Spectra of control experiments of cefuroxime reactivity at 24 h at neutral **(B)** and basic pH **(C)**. Spectra **(D, E)** show the reactivity of cefuroxime with DTT at neutral and basic pH, respectively. The resonance of the β-lactam protons (H6 and H7) was used to monitor the progress of the reaction. Black arrows in (**D**) show the appearance of new chemical shifts for H6 and H7 in the presence of DTT, which indicates the opening of the β-lactam. Blue arrows in (**E**) point out the signals that are decreasing or disappearing as a consequence of a high rate of hydrolysis in the presence of DTT at basic pH (mainly affecting the whole structure except the $R^1$ side chain).
